# Distinct contribution of autoreactive B cell Bruton’s tyrosine kinase signaling to neuroinflammation

**DOI:** 10.64898/2026.04.14.718534

**Authors:** Abraham T. Ogbaslase, Angela S. Archambault, Kia M. Barclay, Benedict E. Ridore, Joshua Amosu, Kaitlyn Ying, Sravanthi Bandla, Alexandria J. Sturtz, Qingyun Li, Peggy L. Kendall, Gregory F. Wu

## Abstract

In multiple sclerosis (MS), autoreactive B cells play a central role in driving CD4 T cell-mediated inflammatory damage to myelin (1). Here we investigated how disrupting Bruton’s tyrosine kinase (BTK) signaling exclusively in B cells shapes the course of experimental autoimmune encephalomyelitis (EAE), a model for MS, through alterations in B cell development and activity. B cell-specific BTK deletion significantly ameliorated both human MOG (hMOG) induced EAE (*p* = 0.0087) as well as spontaneous disease in 2D2^+^IgH^MOG^ mice (*p* = 0.0004). Additionally, MOG-specific cells were found to be more sensitive to loss of BTK than tolerant clones (*p* = 0.0002) and production of anti-MOG immunoglobulins was also found to be diminished (*p* < 0.004) while overall IgG was unchanged (*p* = 0.44). B cells isolated from conditional knockout mice did not upregulate expression of co-stimulatory receptors or MHC II to the same extent as controls when cultured alongside MOG-specific CD4 T cells (*p* < 0.005) and were inferior at driving T cell proliferation (*p* < 0.0001) *in vitro*. Lastly, while BTK deletion diminished the proliferative and survival response of B cells following mitogen stimulation, B cell trafficking to the leptomeninges and organization into ectopic lymphoid tissues (ELTs) in 2D2^+^IgH^MOG^ mice continued unabated. We identified that BTK signaling regulates several features adopted by autoreactive B cells that contribute to EAE pathogenesis. This study provides mechanistic insights into the therapeutic benefits of BTK inhibitors observed in clinical trials exploring BTK as a therapeutic target in the context of MS.

**Significance statement:** Autoreactive B cells contribute to the neuroinflammation that drives multiple sclerosis (MS) and related diseases, yet the molecular mechanisms enabling their pathogenicity remain incompletely understood. This study demonstrates that B cell-specific deletion of Bruton’s tyrosine kinase (BTK) markedly reduces disease severity in two complementary versions of experimental autoimmune encephalomyelitis (EAE), a widely used animal model for MS. Loss of BTK impairs autoreactive B cell survival, antibody production, antigen presentation to encephalitogenic T cells, and T cell activation, while leaving meningeal ectopic lymphoid tissue formation intact. These findings provide direct mechanistic evidence that BTK signaling in B cells promotes neuroinflammatory damage and supports the therapeutic targeting of BTK to limit B cell-driven pathology in MS.

## Introduction

The clinical manifestations of multiple sclerosis (MS) emerge as a result of complex autoimmune targeting of central nervous system (CNS) myelin, most often presenting in a relapsing-remitting course (2, 3). Yet despite treatment with any number of available disease-modifying therapies (DMTs), relapsing MS (RMS) often shifts to a clinical course exhibiting relentless neurologic decline without periods of remission, termed progressive MS (PMS) (4). The cellular and molecular underpinnings of acute and chronic inflammation within the CNS during the course of MS remain unclear.

Increasing evidence points to B cells as key contributors to the propagation of neuroinflammation in MS, through both antibody-independent and antibody-dependent mechanisms (1). The efficacy of anti-CD20 monoclonal antibody therapy for patients with MS strongly underscores the central pathogenic role of B cells (5–9). Models for the role of B cell involvement in MS include versions of experimental autoimmune encephalomyelitis (EAE) in C57BL/6 (B6) mice, such as those involving active immunization with human myelin oligodendrocyte glycoprotein (hMOG), (10) or spontaneous inflammatory demyelination of the CNS resulting from elevated frequencies of myelin specific B and T lymphocytes (11). These animal models have implicated the efficient presentation of antigen by B cells to CD4 T cells via MHCII in perpetuating autoimmune responses in EAE and MS (12). From a pathologic perspective, meningeal inflammation, characterized by aggregates of B cells in the leptomeninges, is often organized into ELT which has been associated with enhanced cortical and spinal cord pathology in PMS (13–15). Additionally, autoreactive B cells can promote neuroinflammation through interactions with microglia. For example, administration of MOG-specific immunoglobulins to mice with EAE promotes Fcγ receptor-mediated proinflammatory activation of microglia (16). Hence, greater understanding of the immune mechanisms that underlie B cell-mediated pathological changes is necessary to develop more effective therapies for MS.

An emerging therapeutic approach for MS involves inhibition of BTK (17). BTK, a Tec family kinase expressed in B cells and most myeloid cells including microglia, has been targeted by a number of small molecule inhibitors for the treatment of autoimmunity (18–22). These small molecule BTK inhibitors (BTKi) are capable of crossing the blood brain barrier (BBB) (17), allowing them to more effectively target microglia as well as peripheral immune cells that have trafficked to the CNS compartment. Inhibition of BTK signaling has been shown to diminish the severity of EAE in mice immunized with whole MOG protein (23). Additionally, treatment of SJL mice with the BTKi evobrutinib reduced the extent of meningeal inflammation following EAE induction (24). While BTK inhibition has promising potential in treating MS (25, 26), off-target effects of BTKi could complicate any therapeutic benefit (21, 26). Furthermore, isolating the effect of BTKi on specific immune populations to determine mechanisms of action is challenging. As such, genetic models readily afford analyses of BTK function in a cell-specific manner. This aids in elucidating the cell-specific contributions to neuroinflammation which is critical not only for mechanistic clarity but also for revealing fundamentally divergent roles of BTK across immune lineages, particularly microglia and B cells. In particular, conditional deletion of BTK allows for evaluating indirect impacts of BTK loss on the phenotype of another population with intact BTK expression. Importantly, immunological changes induced by BTK deletion in mice closely resemble those observed in patients treated with BTKi (17, 27, 28). Accordingly, conditional genetic deletion models not only enable dissection of cell-type-specific contributions of BTK signaling to autoimmune disease progression but also recapitulate key features of pharmacological BTK inhibition in humans.

Herein, we demonstrate that in B cell-dependent EAE, selective deletion of BTK in B cells led to disproportionately impaired development of MOG-specific B cells compared with tolerant counterparts along with a reduction in EAE severity. B cell-specific BTK loss was associated with a reduction in antigen presentation and production of anti-MOG antibodies during EAE. While lymphocyte trafficking to, or organization within, the meninges was not disrupted during EAE in mice lacking BTK in B cells alone, microglial inflammatory responses were constrained. As a result, we have identified a specific role for BTK signaling in B cells orchestrating compartmentalized inflammation during EAE which may underlie improvements in clinical benefit of BTKi in MS.

## Results

### B cell specific deletion of BTK reduces neuroinflammation during EAE

To determine the effect of BTK deletion on the development of EAE, we employed a B cell-dependent model of MS (10) by immunizing global knockout (BTK^KO^) mice with hMOG. We found that BTK^KO^ mice exhibited a milder disease course than WT controls (**Fig. 1A**). To assess the extent to which autoimmune inflammatory damage of the CNS in this model is due B-cell specific BTK signaling, we induced EAE in BTK^ΔCD79^ mice, lacking BTK exclusively in B cells, by immunizing with hMOG. The severity of EAE was dependent on BTK expression by B cells alone (**Fig. 1C**). As BTK plays a key role in B cell development and survival, we were not surprised to see a statistically significant reduction in the percentage (**Fig. 1D**) and number (**Fig. S1**) of B cells in the spleens of BTK^ΔCD79^ mice compared with controls with EAE. However, when using flow cytometry to examine CNS immune cell infiltrates, there was no difference in the frequency (**Fig. 1D**) or number (**Fig. S1**) of B cells within the CNS. Interestingly, B cells that trafficked to the CNS of BTK^ΔCD79^ mice had reduced expression of CD80 and CD86 accompanied by a non-statistically significant reduction in MHC II expression. In comparison, minimal change in expression of these markers was observed in splenic B cells (**Fig. 1E**). Additionally, we found a non-statistically significant reduction in the recruitment of both total CD4 T cells and activated (CD44+) CD4 T cells within the CNS of BTK^ΔCD79^ mice with EAE compared to controls (**Fig 1D, Fig. S1**). As hMOG-induced EAE has been shown to be exacerbated by administration of MOG-specific antibody (29, 30), we evaluated the effect of B cell-specific BTK deletion on the production of anti-MOG antibodies. We found the production of anti-MOG IgG was diminished in BTK^ΔCD79^ mice following EAE induction (**Fig. 1F**) despite similar overall serum IgG levels (**Fig. 1G**). Thus, BTK deletion selectively in B cells diminishes the extent of inflammation during EAE and is associated with impaired B cell antigen presentation.

**Figure 1.**
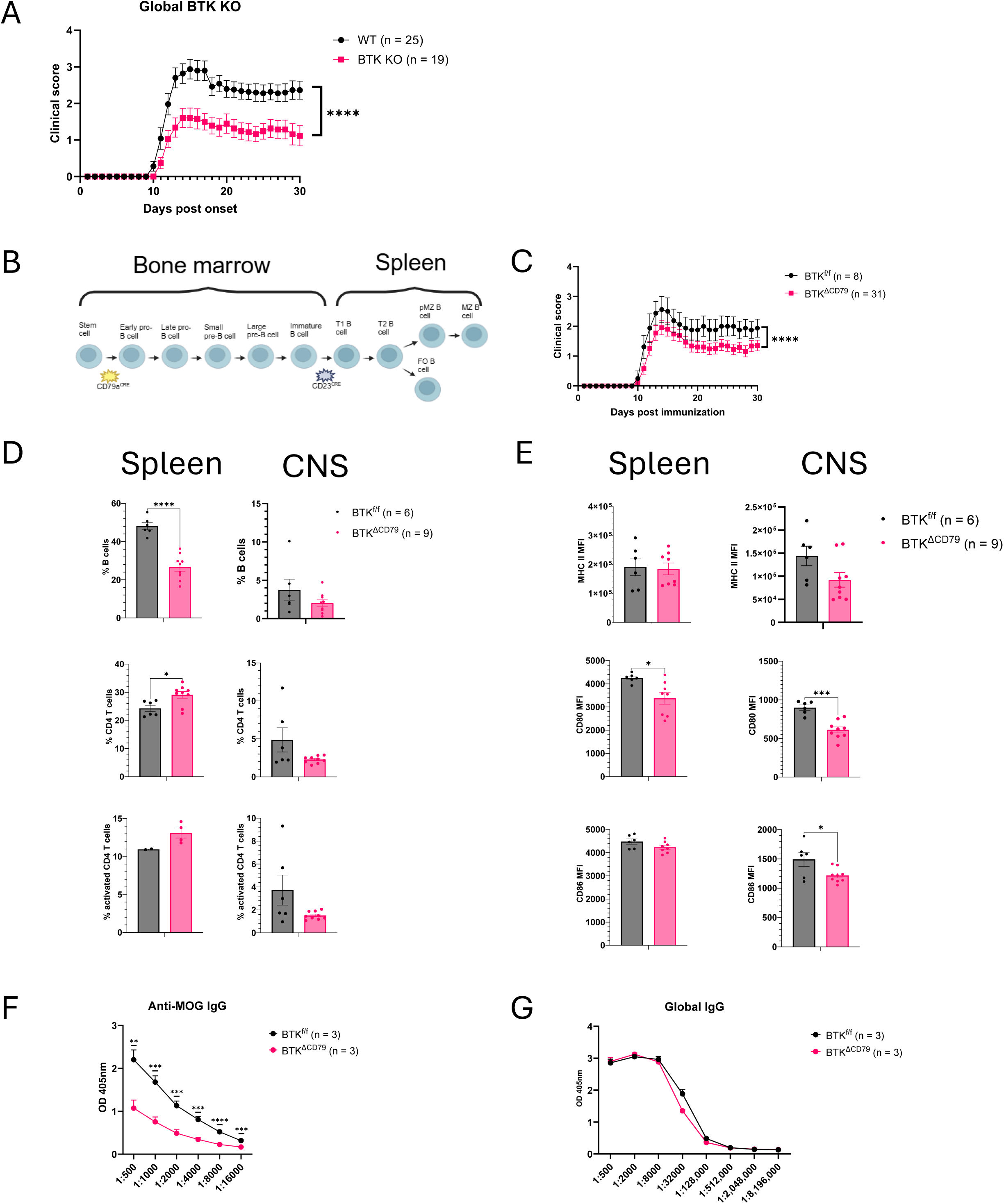
B cell conditional BTK deletion reduces CNS inflammation and EAE disease burden. BTK^-/-^ mice were immunized with hMOG to induce EAE and clinical scores were assessed for 30 days. BTK was floxed out of B cells at the early pro-B cell stage within the bone marrow using a CD79a^CRE^ (B). BTK^f/f^ (black) and BTK^ΔCD79^ (pink) mice were immunized with hMOG to induce EAE and clinical scores were assessed for 30 days (C). Quantification of the frequency of B cells, CD4 T cells and activated CD T cells (CD44+ CD4 T cells) in the spleen (left) and CNS (right, D). B cell expression of MHC II, CD80 and CD86 in the spleen (left) and CNS (right) 30 days post EAE induction in BTK^ΔCD79^ mice (E). Serum anti-MOG IgG (F) and total IgG (G) production in BTK^ΔCD79^ mice 20 days post EAE. P values were determined by Mann-Whitney U test (A, C) or Student’s *t*-test (D-G). Error bars show mean ± SEM; * *p* ≤ 0.05, ** *p* ≤ 0.01, *** *p* ≤ 0.001, **** *p* ≤ 0.0001. (A, C-E) - 3 experiments, (F-G) - 1 experiment.

### The development of MOG-specific B cells and severity of spontaneous EAE depends on B cell BTK expression

With the observation that EAE severity is modulated by B cell expression of BTK, we sought to evaluate the extent to which this phenotype is driven by disruption in B cell development and function following deletion of BTK. In other models of autoimmunity, BTK deletion has been shown to hinder development of autoreactive B cells within the spleen (19, 20). To determine the extent to which BTK contributes to MOG-specific B cell inflammatory responses in neuroinflammation, we crossed BTK^ΔCD79^ mice to 2D2^+^IgH^MOG^ mice (**Fig. 2A**) which express transgenes derived from TCR and B cell IgH loci specific for MOG. As a result, these mice develop EAE spontaneously (11). We observed that BTK deletion in B cells conferred a protective effect against development of spontaneous EAE. 2D2^+^IgH^MOG^ mice lacking B cell expression of BTK exhibited lower disease penetrance as well as reduced disease severity compared to 2D2^+^IgH^MOG^BTK^f/f^ controls (**Fig. 2B**). This phenotype was accompanied by a disruption in B cell development in BTK^ΔCD79^ mice in the 2D2^+^IgH^MOG^ background. Both the frequency and number of B cells were reduced following BTK deletion from B cells in 2D2^+^IgH^MOG^ mice (**Fig. 2C**). Further, the remaining B cells had lower expression of MHC II, CD80, and CD86 (**Fig. 2D, Fig. S2A**). Notably, the development of MOG-specific B cells was disrupted disproportionately in comparison to the rest of the B cell repertoire, with the frequency of MOG-specific B cells reduced from 30.64% ± 1.67% to 17.76% ± 1.26% of B cells following deletion of BTK (*p* = 0.0002; **Fig. 2E**). As steady-state development of B cells is dependent on BTK following emergence from the bone marrow (27, 28, 31), we explored the developmental stage at which BTK deletion disrupts B cell development in the spontaneous model of EAE. We found a developmental blockade of splenic B cells at the transitional 1 (T1) and transitional 2 (T2) stage in BTK^ΔCD79^ mice in the 2D2^+^IgH^MOG^ background (**Fig. 2F, Fig. S2B**). In short, deletion of BTK disproportionately disrupts the production of MOG-specific B cells and confers a protective effect against the development of spontaneous EAE.

**Figure 2.**
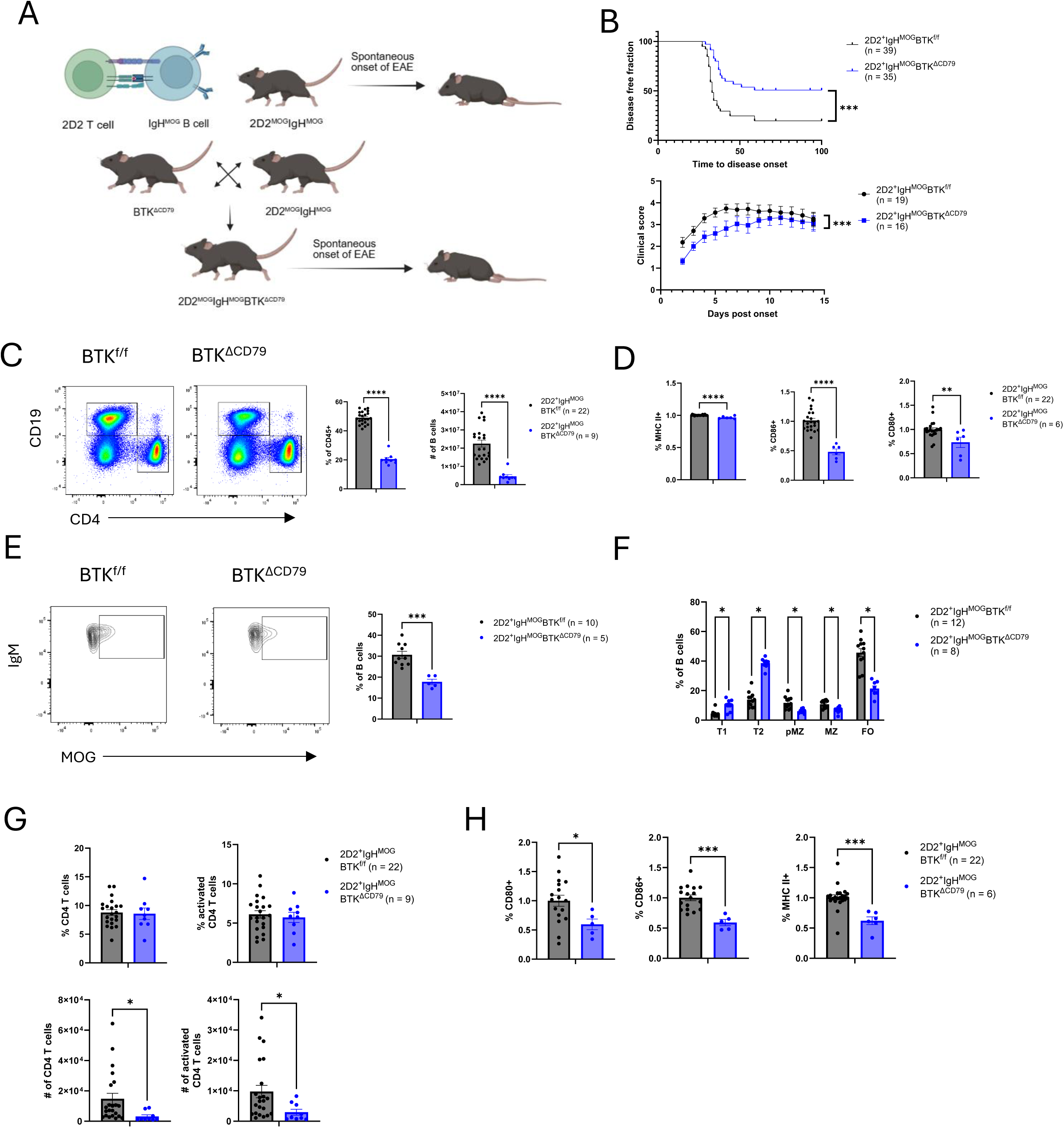
Conditional deletion of BTK in B cells is protective against development of spontaneous EAE. Schematic depicting 2D2^+^IgH^MOG^ mice which express transgenes leading to the development of a high frequency of MOG specific T cells and B cells and resulting in spontaneous onset of EAE (A). BTK^ΔCD79^ mice were crossed to the 2D2^+^IgH^MOG^ background to assess the effect of B-cell specific BTK deletion on disease development in this model. The time to disease onset and disease penetrance (top) as well as disease severity (bottom) was assessed for 2D2^+^IgH^MOG^BTK^ΔCD79^ as well as 2D2^+^IgH^MOG^BTK^f/f^ controls (B). Quantification of the frequency and absolute number of B cells (C) in the spleen of 2D2^+^IgH^MOG^ mice as well as their expression of MHC II, CD80 and CD86 normalized to 2D2^+^IgH^MOG^BTK^f/f^ controls (D). Assessment of the effect of BTK deletion on the production of MOG-specific B cells (E) as well as overall splenic B cell development in 2D2^+^IgH^MOG^ mice (F). Quantification of the frequency (top) and counts (bottom) of CD4 T cells and activated CD4 T cells (CD44+ CD4 T cells) in the brain meninges 14 days post onset of EAE (G). Expression of MHC II and costimulatory receptors on B cells isolated from the brain meninges (H) normalized to 2D2^+^IgH^MOG^BTK^f/f^ controls. P values were determined by Mann-Whitney U test (B) or Student’s *t*-test (C-H). Error bars show mean ± SEM; ns = non-significant, * *p* ≤ 0.05, ** *p* ≤ 0.01, *** *p* ≤ 0.001, **** *p* ≤ 0.0001. (C, D, G, H) - 5 experiments, (E) - 2 experiments, (F) - 4 experiments.

### CD4 T cell infiltration into the brain meninges is diminished by B cell BTK deletion in 2D2^+^IgH^MOG^ mice

One of the unique features accompanying spontaneous inflammatory demyelination in 2D2^+^IgH^MOG^ mice is lymphocytic infiltration of both the brain and spinal cord meninges, which models pathological features observed in PMS (11, 13–15). To evaluate the extent to which the formation of meningeal inflammatory foci is dependent on B cell BTK signaling, we examined 2D2^+^IgH^MOG^ mice 14 days post disease onset and collected meninges from both the brain and spinal cord. We found a reduction in the number of CD4 T cells and activated CD4 T cells within the brain meninges of 2D2^+^IgH^MOG^BTK^ΔCD79^ mice compared with 2D2^+^IgH^MOG^BTK^f/f^ controls (**Fig. 2G**). B cell expression of MHC II, CD80, and CD86 was also reduced within the brain meninges of 2D2^+^IgH^MOG^BTK^ΔCD79^ mice (**Fig. 2H**). Notably, when characterizing the immune response in the spinal cord meninges we identified no difference in T cell infiltration (**Fig. S2C**) or in B cell expression of co-stimulatory receptors or MHC II (**Fig. S2D**). Overall, the protective effect of BTK deletion in B cells during spontaneous inflammatory demyelination was associated with a reduction in T cell recruitment to the brain meninges.

### The formation of meningeal ectopic lymphoid tissues in 2D2^+^IgH^MOG^ mice is not dependent on B cell BTK expression

In PMS, lymphocytes that infiltrate into the meninges often form organized structures resembling ELT that are associated with enhanced cortical and spinal cord pathology (13–15). These aggregations have been described in 2D2^+^IgH^MOG^ mice (11), and we performed IHC to assess the effect of B cell-specific BTK deletion on the formation of ELT within the meninges after disease onset. Meningeal aggregates containing B cells, T cells and in some cases B cells expressing the germinal center marker, GL7, were identified in 2D2^+^IgH^MOG^ mice irrespective of BTK status (**Fig 3A**). We also observed B and T lymphocytes distributed within the meninges in proximity to one another. Interestingly, the numbers of B cells (**Fig. 3B**), T cells (**Fig. 3C**), and GL7+ B cells (**Fig. 3D**) present in meningeal aggregates resembling ELT were unaffected by deletion of BTK within B cells. Thus, the formation of ELT structures in 2D2^+^IgH^MOG^ mice is not dependent on BTK signaling in B cells.

**Figure 3.**
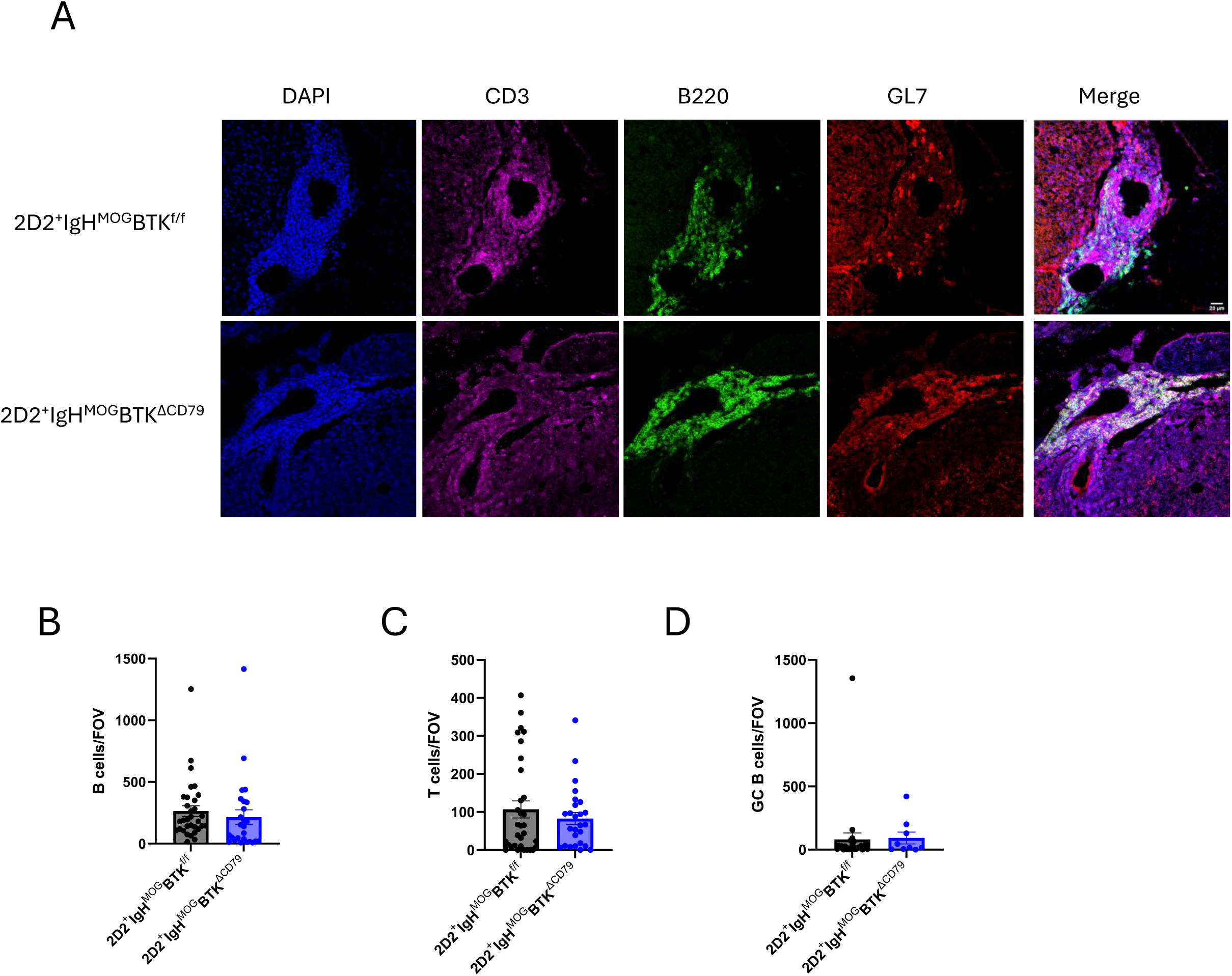
Formation of meningeal ELTs in 2D2^+^IgH^MOG^ mice is not disrupted by deletion of BTK. Representative images of ELTs in the spinal cord meninges (A) were captured and stained for B220, CD3 and GL-7 in both 2D2^+^IgH^MOG^BTK^ΔCD79^ (top) and 2D2-IgH^MOG^BTK^f/f^ (bottom) mice. Quantification of the number of B cells (B), T cells (C) and germinal center B cells (D) per field of view (FOV). Error bars show mean ± SEM.

### Auto-antibody production and APC function by B cells is dependent on BTK signaling during EAE

We sought to characterize the canonical B cell functions being regulated in a BTK dependent manner that contribute to the reduction in disease penetrance seen in 2D2^+^IgH^MOG^BTK^ΔCD79^ mice. Specifically, we investigated how antibody production and antigen presentation capacity of autoreactive B cells in 2D2^+^IgH^MOG^BTK^ΔCD79^ mice were affected by the loss of BTK. Upon examining serum from IgH^MOG^BTK^ΔCD79^ mice, we found that deletion of BTK in B cells significantly reduced the production of anti-MOG IgG. Importantly, overall IgG production was largely unaffected in these mice despite MOG-specific B cells making up a large proportion of the B cell repertoire (**Fig. 4A**). To assess antigen presentation, we isolated B cells from IgH^MOG^ mice and cultured them with CD4 T cells isolated from a naive 2D2 mouse in the presence of 10μg/mL of whole hMOG protein (**Fig. 4B**). After 6 days in culture, we found that the extent of T cell proliferation was significantly reduced in wells that were co-cultured with B cells lacking BTK (**Fig. 4C**). Additionally, B cells derived from BTK^ΔCD79^ mice on the IgH^MOG^ background failed to upregulate CD80, CD86, and MHC II to the same extent as controls (**Fig. 4D & E**). Lastly, we evaluated the ability of B cells to propagate an immune response via cognate interactions with CD4 T cells that have been previously exposed to antigen. To do so, we co-cultured B cells with CD4 T cells isolated from a 2D2 mouse that had been immunized against rhMOG 10 days prior. We found that B cells isolated from mice lacking BTK were inferior at driving proliferation of primed, MOG-specific CD4 T cells compared to controls (**Fig. 4F**). Thus, disruption of BTK signaling impairs the ability of MOG-specific B cells to produce antibodies and dampens their effectiveness as APCs.

**Figure 4.**
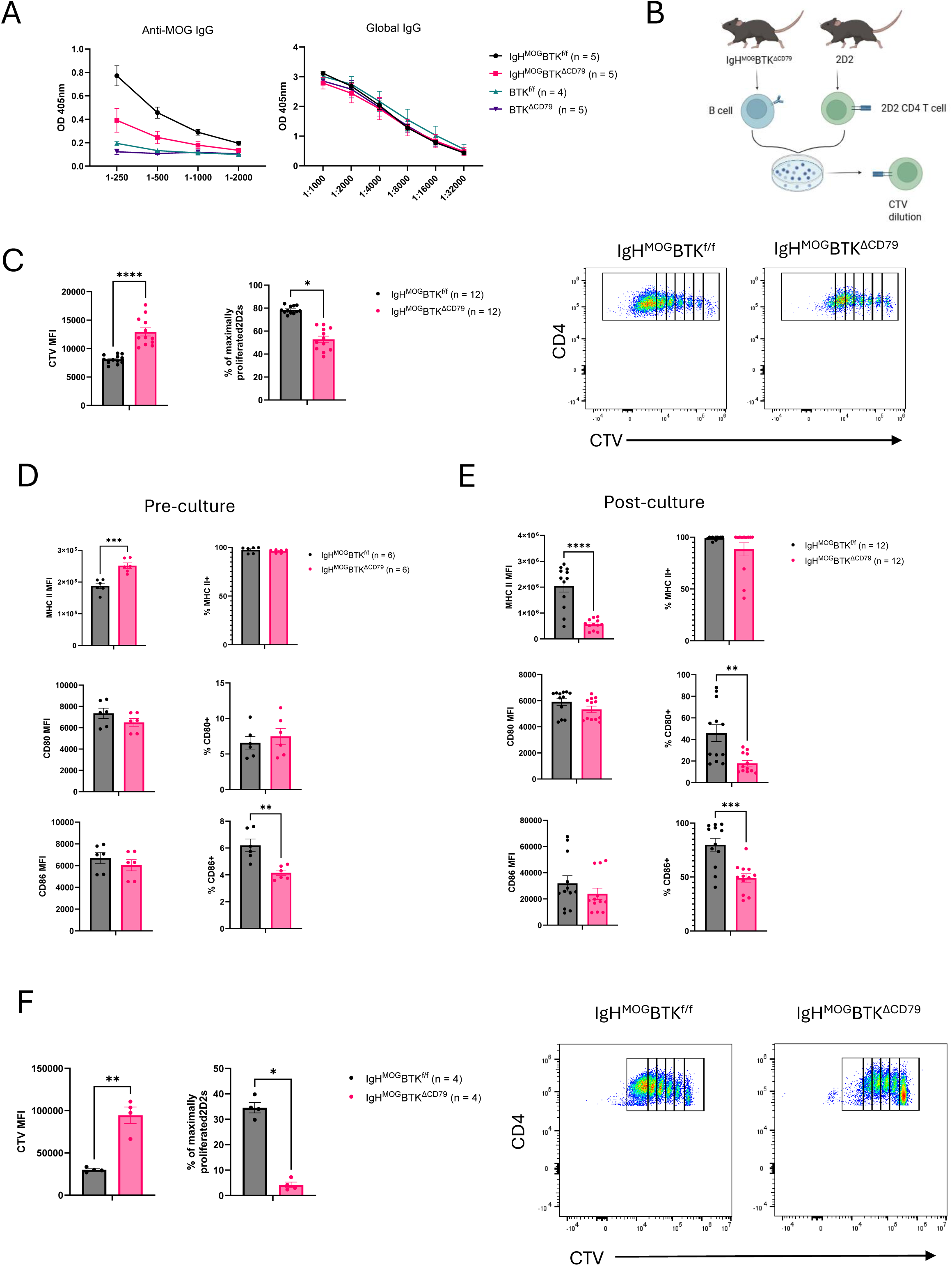
BTK deletion diminishes the production of anti-MOG antibodies and impairs the ability of B cells to drive T cell proliferation. Quantity of anti-MOG IgG and total IgG isolated from the serum of IgH^MOG^BTK^f/f^ (black), IgH^MOG^BTK^ΔCD79^ (pink), BTK^f/f^(teal), or BTK^ΔCD79^ (purple) mice (A). Schematic depicting 2D2 proliferation assay, 2D2 cells were isolated from the spleen of an unimmunized 2D2 mouse, incubated with CTV and co-cultured with B cells isolated from IgH^MOG^ transgenic mice for 6 days (B). The extent of proliferation of 2D2s was quantified by measuring CTV dilution (left) as well as the fraction of cultured 2D2s that had proliferated to the greatest extent (right, C). Quantification of the expression (left) and frequency (right) of MHC II+, CD80+ and CD86+ B cells prior (D) or following (E) culture. Quantification of the extent of CTV dilution in primed 2D2s co-cultured alongside MOG-specific B cells for 4 days (F). P-values were determined by Student’s *t*-test (A, C-F). Error bars show mean ± SEM; ns = non-significant, * *p* ≤ 0.05, ** *p* ≤ 0.01, *** *p* ≤ 0.001, **** *p* ≤ 0.0001. (A, F) - 1 experiment, (C-E) - 4 experiments.

### MOG-specific B cells do not adopt a state of anergy during development

Anergy is a mechanism induced by the immune system to mitigate harm from autoreactive B cells. Anergic B cells are unable to mount a competent immune response upon encountering their antigen and display increased sensitivity to loss of BTK signaling (19, 32–34). However, ectopic expression of MOG in peripheral organs has been shown to completely disrupt development of MOG-specific B cells in the IgH^MOG^ mice (35), suggesting that these B cells may be immunologically ignorant rather than anergic (36). To test the hypothesis that MOG-specific B cells do not adopt a state of anergy during development, we cultured B cells from naive IgH^MOG^ mice in the presence of several mitogens. We found that B cells derived from IgH^MOG^ transgenic mice proliferate to the same degree as those isolated from B6 mice when exposed to anti-CD40 antibodies, IgM cross-linking, or lipopolysaccharide (LPS) (**Fig. 5A**). Additionally, we assessed the viability of IgH^MOG^ transgenic B cells following 3 days in culture. Similarly, we found that there was no difference in the viability of IgH^MOG^ B cells in comparison to those derived from WT mice (**Fig. 5B**). Thus, although spontaneous neuroinflammatory disease in 2D2^+^IgH^MOG^ transgenic mice is sensitive to the loss of BTK, B cells isolated from IgH^MOG^ mice do not adopt a state of anergy.

**Figure 5.**
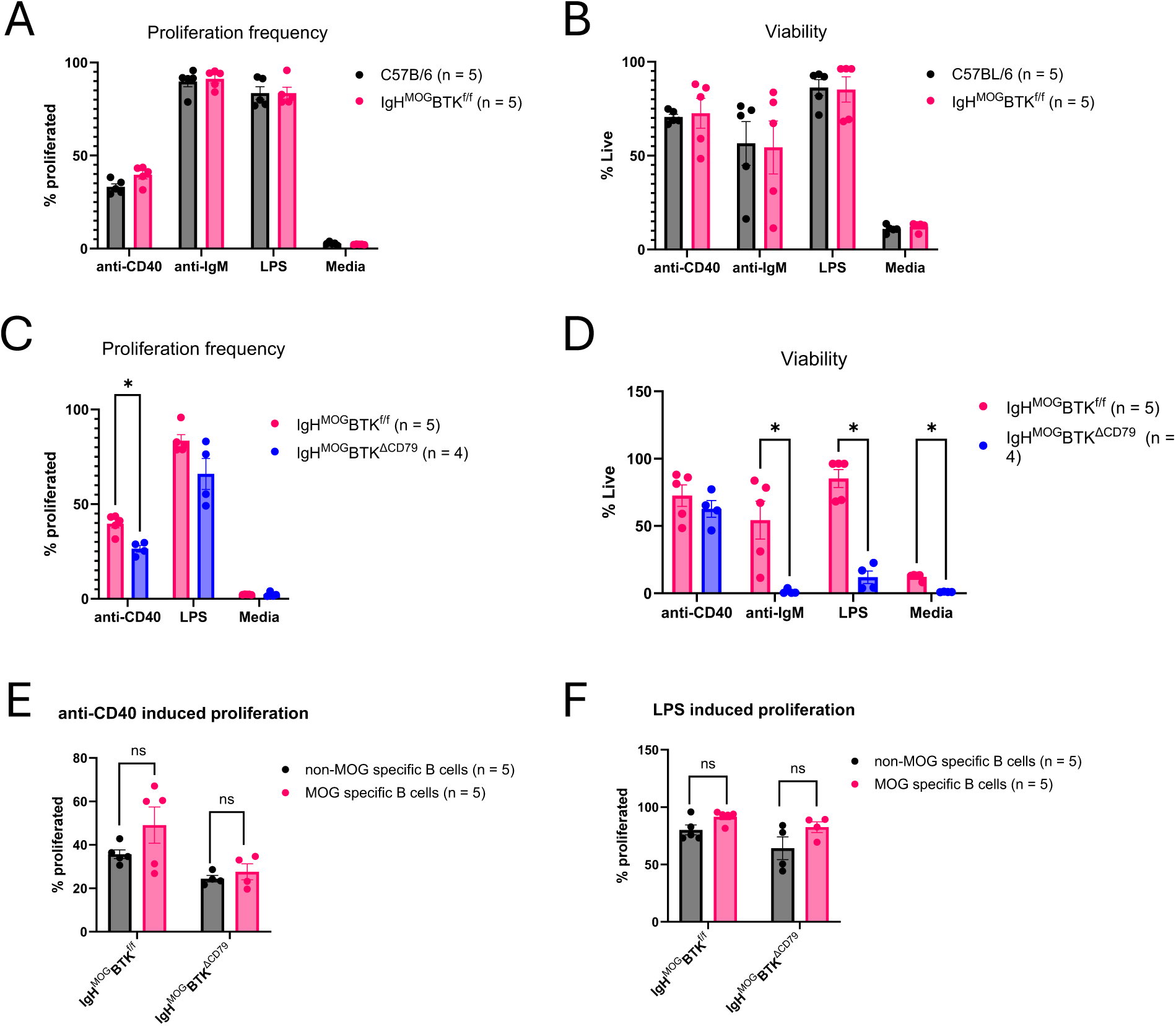
IgH^MOG^ transgenic B cells do not adopt a state of anergy however, proliferation and survival responses to mitogens is compromised by deletion of BTK. B cells were isolated from the spleens of naïve CD57BL/6 (black) and IgH^MOG^BTK^f/f^ (pink) mice and cultured for 3 days in the presence of anti-CD40L (5μg/mL), anti-IgM (10μg/mL), LPS (5μg/mL) or media. Proliferation and survival were assessed by cell trace violet dilution (A) and internalization of zombie NIR viability dye (B) respectively. B cells were also isolated from IgH^MOG^BTK^f/f^ (pink) and IgH^MOG^BTK^ΔCD79^ ^(^blue) to compare how BTK deletion affects proliferative (C) and survival (D) responses. Proliferative responses of MOG-specific B cells (pink) and non-MOG specific B cells (black) to anti-CD40 (E) and LPS (F) stimulation were compared. P-values were determined by Student’s *t*-test (A-F). Error bars show mean ± SEM; ns = non-significant, * *p* ≤ 0.05, ** *p* ≤ 0.01, *** *p* ≤ 0.001, **** *p* ≤ 0.0001. (A-E) - 3 experiments

### Proliferation and survival of IgH^MOG^ transgenic B cells is dependent on BTK signaling

With the observation that B cells isolated from IgH^MOG^ transgenic mice do not exhibit a heightened state of anergy, our next goal was to assess the effect of BTK deletion on the proliferation and survival of B cells derived from IgH^MOG^ mice. Following 3 days in culture, deletion of BTK was associated with impaired proliferative responses of IgH^MOG^ B cells to both anti-C40 and LPS stimulation (**Fig. 5C**). Of note, the viability of B cells following anti-IgM stimulation was too low to assess a proliferative response. Additionally, BTK deletion significantly impaired B cell survival in response to LPS stimulation but not anti-CD40 stimulation (**Fig. 5D**). Lastly, we characterized the proliferative response based on the specificity of the B cell receptor. We found that proliferation of MOG-specific B cells did not differ significantly from that of non-MOG-specific B cells (**Fig. 5E & F**). Thus, the survival and proliferative responses of autoreactive B cells derived from IgH^MOG^ mice are diminished following deletion of BTK irrespective of the B cell antigen specificity.

### BTK deletion later in B cell development confers protective effect against development of EAE

The extent to which B cell dependence on BTK drives neuroinflammation in EAE is related to disruption of B cell development versus B cell activation remains unclear. To better understand how disturbing B cell activation following BTK deletion contributes to this protective phenotype in the context of EAE, we crossed BTK^f/f^ mice to mice expressing Cre recombinase under the CD23 promoter (BTK^ΔCD23^), a gene which is expressed in late-stage B cell development (**Fig. 1B**). We found that BTK^ΔCD23^ mice experience a milder disease course than BTK^f/f^ controls (**Fig. 6A**). This protective phenotype was observed despite the effect of BTK deletion on B cell development being muted in this mouse strain (**Fig. S3B-D**). Interestingly, overall antibody and anti-MOG antibody production were both diminished in BTK^ΔCD23^ mice following EAE induction, with MOG-specific IgG most affected (**Fig. 6B**). Thus, the protective phenotype mediated by B cell-specific BTK deletion in the context of EAE is not dependent on the timing of B cell developmental disruption.

**Figure 6.**
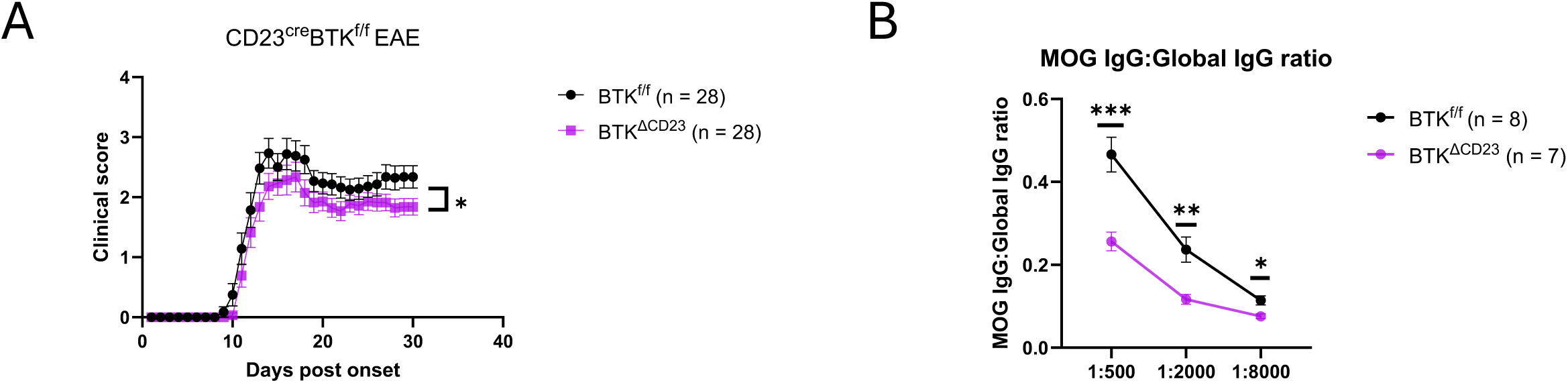
Deletion of BTK within developmentally mature B cells confers a protective effect against development of EAE and ameliorates neuroinflammation. BTK^f/f^ (black) and BTK^ΔCD23^ (purple) mice were immunized with hMOG to induce EAE and clinical scores were assessed for 30 days (A). Quantification of anti-hMOG IgG and total IgG (B) isolated from the serum of BTK^f/f^ (black) or BTK^ΔCD23^ (purple) 30 days post immunization. P values were determined by Mann-Whitney U test (A) or Student’s *t*-test (B). Error bars show mean ± SEM; ns = non-significant, * *p* ≤ 0.05, ** *p* ≤ 0.01, *** *p* ≤ 0.001, **** *p* ≤ 0.0001. (A) - 4 experiments, (B) 1 experiment

## Discussion

Herein, we have shown that B cell conditional BTK deletion diminishes the antigen presentation capacity of B cells. In the absence of BTK, B cells do not efficiently promote proliferation of MOG-specific CD4 T cells *in vitro* nor drive neuroinflammation *in vivo.* Despite MOG-reactive B cells failing to adopt features of anergy, development of autoreactive B cells was found to be more susceptible to loss of BTK than non-self-reactive counterparts. This was characterized *in vivo* through increased dropout of MOG-specific B cells following BTK deletion and by a diminished proliferative and survival response following mitogen stimulation *in vitro*. In summary, our results provide mechanistic insight into the potential of BTK inhibition to selectively impair pathogenic B cell function and limit B cell-dependent neuroinflammation in MS.

The animal models of MS used in this study were purposely selected to highlight the contribution of B cells to neuroinflammatory disease development and propagation. Immunization of B6 mice with hMOG induces a B cell-dependent disease course (10). Similarly, IgH^MOG^ mice greatly amplify the frequency of spontaneous disease in the 2D2 background (11). The B cell dependency of these models allowed us to explore the contribution of unique B cell characteristics, such as antigen presentation, to the pathophysiology of MS. Further, the central role of MOG-reactive BCRs in driving B cell activation in both hMOG immunization and spontaneous versions of EAE narrows the focus on BTK, a kinase critical for transducing BCR signals into pro-inflammatory and pathogenic B cell responses. These models, therefore, facilitated our observations that in addition to changes in the ability of B cells to present antigen, we found BTK deletion disproportionately reduced the production of anti-MOG antibodies. Individually, both B cell antigen presentation (12) and administration of anti-MOG antibodies (29) have been shown to contribute to the development of EAE and in this study we have established that both of these features are regulated in a BTK dependent manner. Ultimately, we did not use more commonly employed disease models in which B cells are dispensable in order to avoid masking the consequences of B cell-conditional BTK deletion.

We established that autoreactive B cells specific for MOG failed to adopt a state of anergy during development. This is in stark contrast to insulin-specific B cells studied in murine models of diabetes (19, 34). The divergence between autoreactive B cell populations specific for MOG and insulin may be due to differences in the structure and location of the antigen to which B cells are responding. Unlike insulin, which is a small, soluble protein, MOG is relatively large and expressed on the cell membrane (37). As a result, MOG has a greater ability to induce BCR cross-linking upon recognition by B cells and proximal BCR signaling which may underlie the B cell developmental differences observed in these models of tissue-specific autoimmunity (33). Additionally, unlike insulin, which is expressed in the periphery, expression of MOG is restricted to the CNS. A lack of exposure to MOG in the periphery may result in MOG-specific B cells not being recognized as autoreactive during development and may encourage B cells towards immunological ignorance rather than anergy. The observation that BTK deletion impairs the development and activity of functionally competent autoreactive B cells supports the idea that BTKi will effectively dampen the pathogenicity of autoreactive B cells in the context of MS.

Along with the lack of anergic responses by MOG-specific B cells, we found that BTK deletion significantly impairs the survival response of IgH^MOG^ derived B cells upon stimulation with LPS or anti-IgM, but not following exposure to anti-CD40. At this point we hypothesize that this impaired viability is due to disruption of NFkB signaling, which promotes B cell survival, following BTK deletion. TLR signaling has been shown to supplement proximal BCR signaling through the adaptor protein MyD88 (38). Deletion of BTK may disturb the ability of these mitogens to induce NFkB signaling resulting in B cell death. Interestingly, CD40 signaling also promotes B cell survival through NFkB signaling; however, NFkB activation in this pathway is dependent on E3 ubiquitin ligase activity rather than tyrosine kinase signaling (39). This difference may account for the competent survival response of B cells in the face of BTK deletion.

The meninges of 2D2^+^IgH^MOG^ mice features ELTs which are characterized by lymphocyte aggregation and the presence of GL7+ B cells. These parallels to germinal centers present within secondary lymphoid organs suggest that there is a unique immune response occurring within the meninges of 2D2^+^IgH^MOG^ mice. Similar structures are also found within the meninges of PMS patients where there presence is associated with increased disease severity (13–15). However, the protective effect of B cell conditional BTK deletion does not seem to be related to the presence of these structures since we were unable to detect differences in abundance when in 2D2^+^IgH^MOG^BTK^ΔCD79^ mice developed neurologic disease. It is possible that a certain threshold of meningeal inflammation needs to be crossed before a cascade of events, independent of B cell BTK expression, unfolds. Nevertheless, it remains to be seen whether BTKi can result in a reduction in mELT in MS.

Additionally, while B-cell conditional BTK deletion significantly ameliorates the disease course following hMOG induced EAE, we found that this protective phenotype is exaggerated following global BTK deletion. This suggests that BTK expression in myeloid cells, such as microglia meaningfully contributes to disease pathogenesis in this model of EAE. Not only is microgliosis a prominent feature observed in PMS, microglia isolated from CNS tissue of ALS patients have been shown to upregulate expression of BTK and treatment with BTKi has been shown to reduce the extent of neuronal apoptosis induced upon exposure to microglial conditioned media (40, 41). In the future, we are particularly interested in exploring the effect of disrupting microglial BTK signaling on neurological disease progression.

A limitation of this study is that we did not assess the effect of administering BTKi on B cell phenotype or susceptibility to EAE. We intentionally developed our experiments around conditional genetic systems to minimize the off-target effects of BTKi. These side-effects, including liver toxicity, have complicated clinical trials evaluating the efficacy of BTKi for MS (26). However, it is possible that some of the clinical benefits of BTKi are due to targeting kinases other than BTK. Evaluating the difference between BTKi treatment and our conditional knockout models would enable us to better discern how the non-specific effects may contribute to either the benefit or detriment of inhibitors’ clinical outcomes. Additionally, we only assessed the effect of constitutive BTK deletion on the development of EAE. Use of a tamoxifen-inducible system would have allowed us to evaluate the effect of therapeutic BTK deletion on disease outcomes which could serve as an alternative reflection of various BTKi treatment being tested currently in MS.

Our findings carry real-world implications for the translatability of BTKi for MS. Given the involvement of B cell signaling in the development of MS (5, 7, 42), the observation that BTK deletion impairs the development and activity of otherwise functionally competent B cells suggests that BTKi will have the ability to compromise the pathogenicity of B cells in MS patients. The amelioration of disease in BTK^ΔCD23^ mice, in which BTK deletion occurs when cells are traversing the early checkpoints of peripheral B cell tolerance at the T1 stage and beyond, raises intriguing questions about the mechanisms by which BTK shapes the autoreactive B cell repertoire. While this model does not cleanly isolate B cell activation from development, it does shift the developmental window during which BTK is required, with the surviving B cells still subject to the full gauntlet of transitional tolerance. Nonetheless, the protection of disease observed in BTK^ΔCD23^ mice, despite a comparatively modest effect on overall B cell numbers, suggests that BTK-dependent processes operating in transitional and mature B cells beyond the earliest developmental checkpoints contribute to B cell-dependent EAE. Our collection of results support the concept that autoreactive B cells escaping early tolerance and persisting within the mature B cell repertoire remain disproportionately reliant on BTK for survival and activation signals. Hence, autoreactive B cells represent a selectively vulnerable subset which are exploitable for BTKi based on the intrinsic BTK dependency in the treatment of MS.

## Materials and Methods

### Mice

B6 mice were bred and housed in a single pathogen-free housing facility at Washington University School of Medicine and fed mouse chow and water ad libitum. Global BTK^KO^ mice were obtained from Dr. Peggy Kendall at Washington University School of Medicine. Exons 13 and 14 of BTK were replaced with neomycin in BTK^KO^ mice, resulting in deficits in B cell development and activation (43). B cell conditional BTK knockout mice were made by crossing mice expressing a CD79a^cre^ line (44) to BTK^f/f^ mice (18) (BTK^ΔCD79^; **Fig. 1B**). Additionally, BTK^f/f^ mice were crossed to a CD23^cre^ line (BTK^ΔCD23^ mice), leading to BTK deletion in transitional and mature B cells. Cre activity can also be detected in a small subset of immature B cells in BTK^ΔCD23^ mice but is absent in pre- and pro-B cells (45), thus allowing us to primarily focus on the effect of BTK deletion on B cell activation rather than development. Male and female mice between 6 and 10 weeks of age were used to induce active EAE. MOG peptide T cell receptor transgenic mice were bred to MOG-specific B cell receptor (BCR) knock-in (IgH^MOG^) mice to generate 2D2^+^IgH^MOG^ transgenic mice which develop EAE spontaneously (11). These mice were subsequently crossed to BTK^ΔCD79^ mice and evaluated for 14 days post-onset of EAE to characterize the effect of B cell-specific BTK deletion on disease course.

### Induction of active EAE

EAE was induced by subcutaneously injecting 150 μg of recombinant human MOG (rhMOG) protein emulsified in complete Freund’s adjuvant (CFA; Sigma) containing 500 μg H37RA (Difco, Franklin Lakes, NJ). Additionally, mice were intraperitoneally injected with 400 ng of pertussis toxin in 200 µL of PBS on days 0 and 2 post-immunization. EAE was scored using the following rubric: 1 – loss of tail tone, 2 – impaired righting reflex, 3 – partial hindlimb paralysis, 4 – complete hindlimb paralysis, 5 – death.

### Flow cytometry

Prior to perfusion, spleens were harvested from mice and processed through a 40 μm filter into single cell suspensions. These suspensions were then treated with ACK erythrocyte lysis buffer. Brain and spinal cord tissue were collected from mice following perfusion with 30 mL of ice-cold PBS and homogenized through a 40 μm filter into single-cell suspensions. CNS cells were purified by centrifugation for 30 minutes at 300g in a solution consisting of 30% Percoll (Cytiva, Washington, D.C.) and 10% 10X Earle’s Balanced Salt Solution (EBSS, Sigma, St. Louis, MO). The following antibodies were purchased from Biolegend (San Diego, CA): IgM – BV711, IgMa – PerCPCy5.5, IgD – APC/Cy7, CXCR5 – AF647, CD19 – BV750, CD69 – Spark NIR 685, CD44 – BV650, CD62L – BV785, TCRb – BV570, CD4 – BV605, CD8 – Spark YG 593, CD80 – PECy5, CD86 – PE Dazzle 594, MHC II – PE Fire 810, Ly6G – BV421, CD11c – PE Fire 700, CD93 – APC, CD23 – FITC, CD21 – PECy7, SA – BV421, SA – BV510, Zombie Red. The following antibodies were purchased from ThermoFisher (Waltham, MA): CD25 – BUV737, CD11b – BV480, Ly6c – eFluor 450, CD45.2 - AF532. Samples were run on a Cytek Aurora 3 Laser spectral flow cytometer and analyzed with FlowJo (BD Biosciences) software.

### Anti-MOG antibodies

Anti-MOG antibody production was quantified by coating a 96 well ELISA plate overnight with 100 μL of a 10μg/mL solution of rhMOG. The following day, wells were incubated with 200 μL of a blocking solution (PBS + 1% BSA) prior to adding serial dilutions of serum samples to each well overnight. The next day, wells were incubated with alkaline phosphatase-conjugated goat anti-mouse IgG secondary antibody (Jackson, 115-056-072, 1:5000 dilution) followed by a p-Nitrophenyl phosphate (PNPP) solution (Sigma, N1891). After 30 minutes, absorbance at 405nm was measured.

### Meningeal isolation

Fourteen days post-EAE onset, dura mater was collected from the brain and spinal cord of 2D2^+^IgH^MOG^ mice (46). To do so, mice were first perfused with 30 mL of cold PBS and meningeal tissue was isolated from the skull and spinal column. Isolated tissue was then chemically digested in digest buffer (HBSS + 0.8mg/ml dispase II + 0.2mg/mL collagenase) for 25 minutes at 37°C. Following this incubation, samples were mechanically digested and filtered into FACS tubes for staining and analysis.

### Histology

Mice were sacrificed and perfused with 30 mL of ice-cold PBS. Brain and spinal cords were removed within the skull and spinal column, respectively, and fixed in 4% paraformaldehyde (PFA) overnight. Tissues were then decalcified in a 6% trichloroacetic acid solution for 5 days before dehydrating in 30% sucrose overnight or longer. Tissues were then paraffin-embedded and sectioned at 10 μm, with sections stained for B cells (primary antibody: Abcam, Ab245235, rabbit anti-mouse CD19, secondary antibody: Invitrogen, A11034, goat anti-rabbit AF488), T cells (primary antibody: Biolegend, 100302, Armenian hamster anti-mouse CD3, secondary antibody: Invitrogen, A78967, goat anti-Armenian hamster AF647), germinal center B cells (primary antibody: Invitrogen, 14-5902-82, rat anti-mouse GL7, secondary antibody: Invitrogen, A21434, goat anti-rat AF555) and/or microglia (primary antibody: FujiFilm, 019-1974, rabbit anti-mouse Iba1, secondary antibody: Invitrogen, A11034, goat anti-rabbit AF488). Slides were examined by confocal microscopy on the Nikon A1R at the Washington University Center for Cellular Imaging (WUCCI) and analyzed using ImageJ.

### T cell proliferation assay

To assess proliferation, 2D2 T cells were purified from unimmunized 2D2 mice and cultured with B cells isolated from naive IgH^MOG^ mice. Both B cells and T cells were isolated through positive selection using CD19 (Miltenyi, 130-121-301) and CD4 microbeads (Miltenyi, 130-117-043), respectively. Samples were then passed through a magnetic column (Miltenyi, 130-042-401) to separate labelled cells. Each well consisted of 100,000 T cells with 100,000 MOG-specific B cells in CTS Optimizer T cell expansion media (Thermofisher, A1048501) reinforced with 10 IU/mL of human IL-2 and 10 μg/mL of rhMOG protein. Cells remained in culture for 6 days with fresh media being added on day 3. Proliferation was evaluated by measuring the extent of cell trace violet (CTV) dilution. To assess proliferation of primed 2D2 T cells the same protocol was followed, but T cells were isolated from 2D2 mice that had been immunized with rhMOG 10 days prior and cells remained in culture for only 2 days. Cells cultured with 10 μg/mL of anti-CD3 or with no MOG were used as positive and negative controls, respectively.

### B cell proliferation assay

To assess the functional state of B cells isolated from IgH^MOG^ mice, we cultured 1×10^6^ B cells in a 24 well plate for 3 days. B cells proliferation was induced by adding one of the following mitogens: 5 μg/mL of anti-CD40, 10 μg/mL of anti-IgM, or 5 μg/mL of LPS. Proliferation was assessed by quantifying the extent of CTV dilution.

---

**Supplemental Figure 1.**
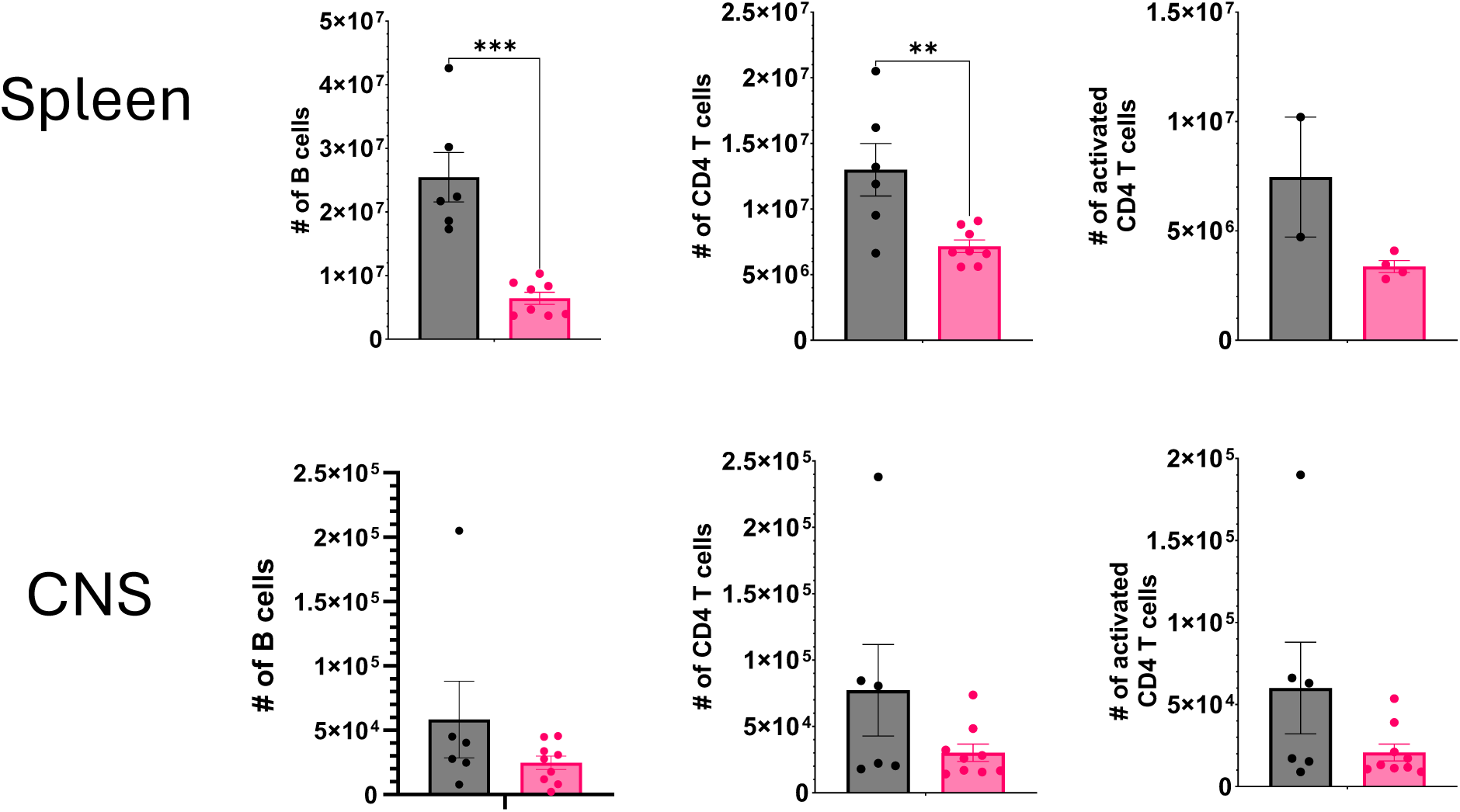
B cell conditional BTK deletion reduces the number of lymphocytes in the periphery following EAE induction. Quantification of the number of B cells, CD4 T cells and activated CD T cells (CD44+ CD4 T cells) in the spleen (left) and CNS (right) 30 days following induction of EAE (A). P values were determined by Mann-Whitney U test. Error bars show mean ± SEM; ns = non-significant, * *p* ≤ 0.05, ** *p* ≤ 0.01, *** *p* ≤ 0.001, **** *p* ≤ 0.0001. 3 independent experiments.

**Supplemental Figure 2.**
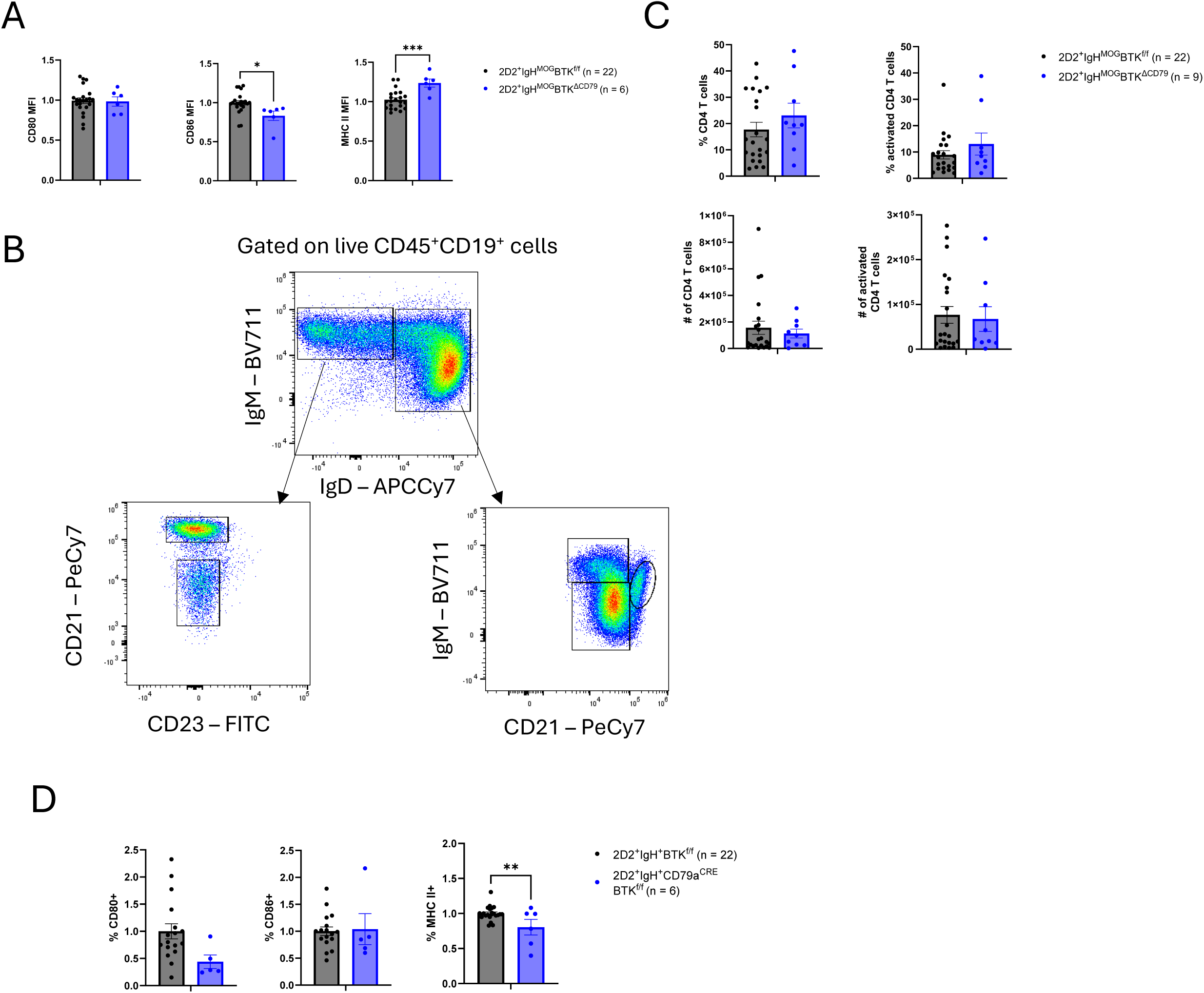
Effect of B cell conditional BTK deletion on lymphocyte development and activation in 2D2^+^IgH^MOG^ mice. Splenic B cell expression of MHC II, CD80 and CD86 normalized to 2D2^+^IgH^MOG^BTK^f/f^ controls (A). Gating strategy for identifying splenic B cell populations in 2D2^+^IgH^MOG^ mice (B). Quantification of the frequency (top) and counts (bottom) of CD4 T cells (left) and activated CD4 T cells (CD44+ CD4 T cells, right) in the spinal cord meninges 14 days post onset of EAE (C). Expression of MHC II and costimulatory receptors on B cells isolated from the spinal cord meninges normalized to 2D2^+^IgH^MOG^BTK^f/f^ controls (D). P values were determined by Student’s *t*-test. Error bars show mean ± SEM; ns = non-significant, * *p* ≤ 0.05, ** *p* ≤ 0.01, *** *p* ≤ 0.001, **** *p* ≤ 0.0001. (A, C, D) - 5 experiments, (B) – 4 experiments.

**Supplemental Figure 3.**
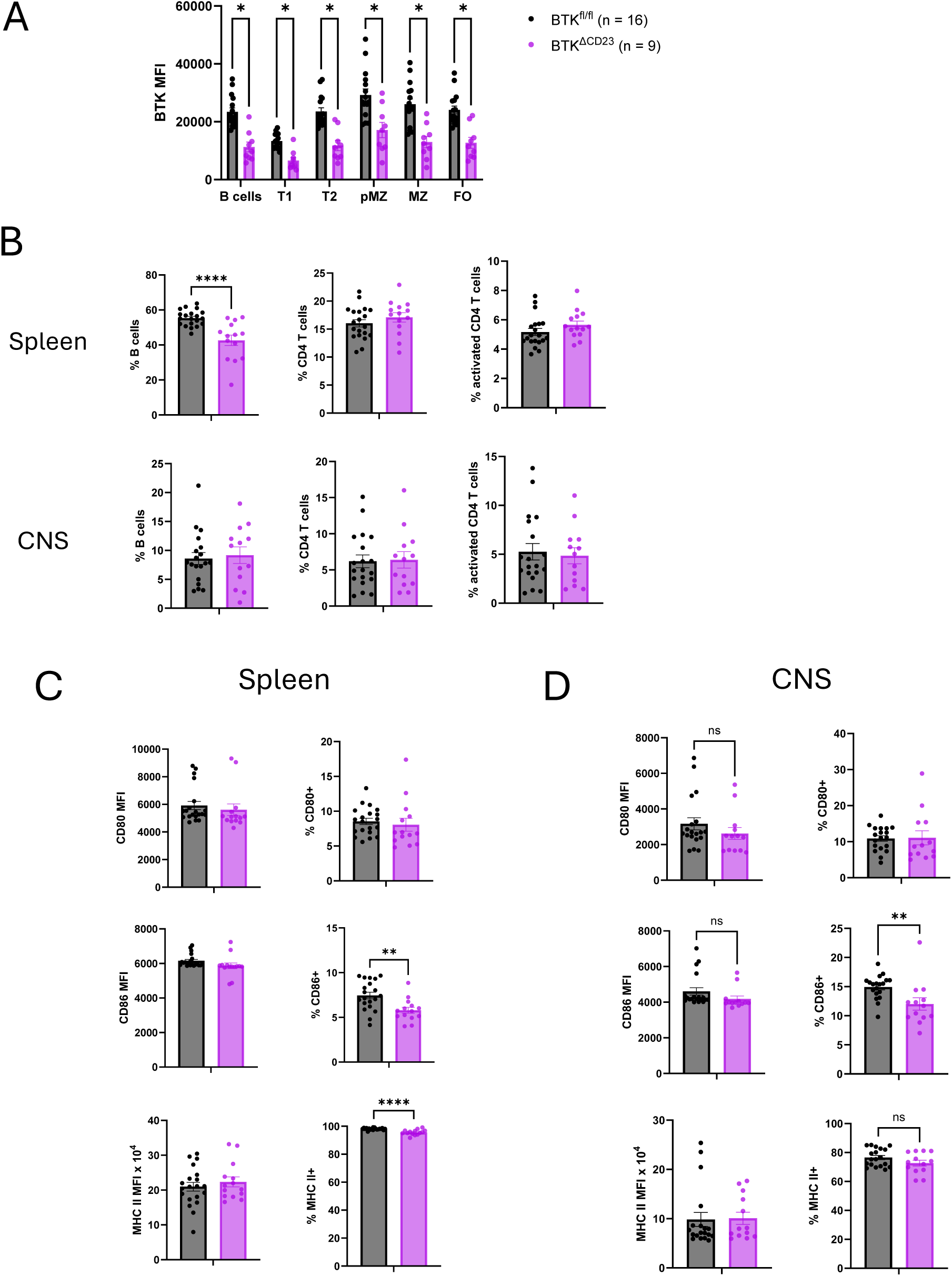
Immune infiltrate in BTK^ΔCD23^ mice following induction of EAE. Quantification of BTK expression in splenic B cell populations 30 days post induction of EAE (A). Quantification of the frequency of B cells, CD4 T cells and activated (CD44^+^) CD4 T cells in the spleen (top) and CNS (bottom, B). B cell expression of MHC II, CD80, and CD86 in the spleen (C) and CNS (D) 30 days post EAE induction. P values were determined by Student’s *t*-test. Error bars show mean ± SEM; ns = non-significant, * *p* ≤ 0.05, ** *p* ≤ 0.01, *** *p* ≤ 0.001, **** *p* ≤ 0.0001. (A-D) – 3 experiments.

